# Divergence and parallelism in two tropical drosophilids simultaneously invading a desert environment

**DOI:** 10.1101/2024.10.02.616270

**Authors:** Ahmed M. El-Sabrout, Céline Moreno, Mélody Temperville, Erina A. Ferreira, David Ogereau, Issa Mze Hassani, Héloïse Bastide, Amira Y. Abou-Youssef, Amir Yassin

## Abstract

Invasive species have once been called a ‘grand experiment in evolution’ but natural replicates of such experiments are often scarce. In particular, whether the rapid adaptation to the new environment involves genetic predisposition in the ancestral range or mainly occurs via post-introductory selection on a genetically variable propagule remains unknown. Here, we investigate the parallel adaptation of two drosophilid species of the genus *Zaprionus*, *Z. indianus* (the African fig fly) and *Z. tuberculatus*, to contrasting agrarian and desert environments following their introduction in Egypt during the last four decades. Field collection unraveled distinct spatial distribution of the two species. Population genomics analyses showed correlated differentiation levels at orthologous genes before and after introduction in both species. Nonetheless, phenotypic analyses revealed distinct fruit preference and desiccation resistance between both species as well as between introduced and native *Z. tuberculatus* populations. Hence, despite signals of genomic parallelism, ecological divergence between the two species likely facilitates their co-existence in the introduced regions. Our results provide a significant step towards understanding the mechanisms underlying the simultaneous invasive success of both species, which have also recently invaded the Americas and Europe, and of which one at least is a notorious pest.

## Introduction

Invasive species are a major threat to biodiversity (Capinha et al. 2015; Bellard et al. 2016; Bradshaw et al. 2016; Early et al. 2016). With the accelerating rate of human-mediated habitat homogenization and climate warming, species that are better adapted to human settlements and transportation and/or hotter and drier climates are expected to overwhelm many native environments (Capinha et al. 2015; Fournier et al. 2019).

Understanding how such species rapidly adapt to the new environments to which they are introduced is an important challenge for invasion genetics. The development of recent genomic tools provides a valuable opportunity to address questions like the routes and number of introduction events, the effect of the introduction bottlenecks on genetic variation, as well as the relative role of phenotypic plasticity and/or ancestral genetic variation (*i.e.* genetic predisposition) to adaptation to the new environment (North et al. 2021). From an evolutionary biology point of view, invasive species constitute a ‘natural experiment’ to track rapid adaptation in real-time. In some cases, multiple introductory events of a single species in different geographical localities with similar habitats constituted unique natural replicates (Ayala et al. 1989; Bergland et al. 2016; Calfee et al. 2020; Stern and Lee 2020; van Boheemen and Hodgins 2020; Louis et al. 2021; Pélissié et al. 2022; Stuart et al. 2022). However, because most of these parallel introduction cases usually share significant proportions of variation from the ancestral range, it remains unclear whether parallelism could also underlie invasion success in species belonging to distinct evolutionary lineages.

The expansion of the drosophilid species *Zaprionus indianus* from its native range in Tropical Africa to the Americas up to Hawaii islands in less than 20 years is a spectacular example of the rapid adaptive success of a human-associated biological invasion (Gibert et al. 2016; Bragard et al. 2022). The species was first reported in southern Brazil in 1998 where it was responsible for the loss of nearly 40% of fig fruits production (Bragard et al. 2022). It was therefore called the ‘African fig fly’, although it has been bred from more than 80 plants including some native Brazilian plants and it is thought to be a potential pest for multiple soft-skinned fruit crops (Leão and Tidon 2004; Yassin and David 2010; Bragard et al. 2022). The species’ Latin name is also ill- coined, referring to its first description from India where it has likely been introduced in the 1960s. From this independent, older introduction the species has slowly yet steadily expanded in the Palearctic region (the Arabian Peninsula and around the Mediterranean basin) in the west and up to Bangladesh in the east (Yassin et al. 2008b; EFSA Panel on PlantHealth (PLH) 2022). Parallel to these invasions, another Afrotropical species of the genus *Zaprionus*, *Z. tuberculatus*, has also been introduced in the Mediterranean basin (Cyprus, Egypt, Israel and Malta) in the early 1980s (Chassagnard and Kraaijeveld 1991), and expanded northward since the 2010s at a slower pace toward Turkey, southern Europe and Russia (Oboña et al. 2019; Özbek Çatal et al. 2021). In 2021, it was recorded for the first time in Brazil indicating its likely future spread across the Americas (Cavalcanti et al. 2022). Although *Z. tuberculatus* has almost the same polyphagous fruit-breeding habit as *Z. indianus*, its potential as an agricultural pest remains to be evaluated (Cavalcanti et al. 2022; Viana et al. 2024).

The simultaneous invasion of the Palearctic region by the two *Zaprionus* species hence represents a unique opportunity to investigate the genomic basis of invasion success. The two species are distantly related; each belongs to distinct species groups that have diverged 7-14 million years (myr) ago (Yassin et al. 2008a; Suvorov et al. 2021). Therefore, any genetic parallelism between the two species is more likely due to selection working on independent mutations rather than on a shared variant. Here, we conduct a comparative population genomics analysis in both species to investigate possible signals of divergence parallelism occurring between agrarian vs. desert populations in Egypt and Egyptian vs. Afrotropical populations. We found that despite signals of parallelism in genome-wide differentiation patterns before and after the introduction of the two species in Egypt, ecological, behavioral and physiological differences likely facilitate their coexistence in the introduced regions.

## Materials and Methods

### Fly collection

Flies were collected in Egypt in November 2019 from two localities: Wadi El Natrun (WN, ∼120 km southwest of Alexandria), a desert reclaimed farm producing citrus and date palms, and Damanhur (DM, ∼70 km southeast of Alexandria), an agrarian area in the Nile Delta (Figure 1A). In the first locality, fermented banana traps were hanged on branches of citrus trees, whereas in the second locality four collection sites were chosen: a guava orchard (D30), a house garden surrounded by date palm trees (WG), a house roof (HH) and a beet field with no fruit production (GH) (Figure 1B). Flies were immediately placed in absolute ethanol and then taxonomically sorted under a binocular microscope, using published taxonomic keys (Yassin and David 2010). The two *Zaprionus* species are readily distinguishable and cannot be confused on the basis of the morphology of the forefemora in both sexes, bearing a series of spines in *Z. indianus* and a single long bristle bore on a protruding wart in *Z. tuberculatus*.

**Figure 1.**
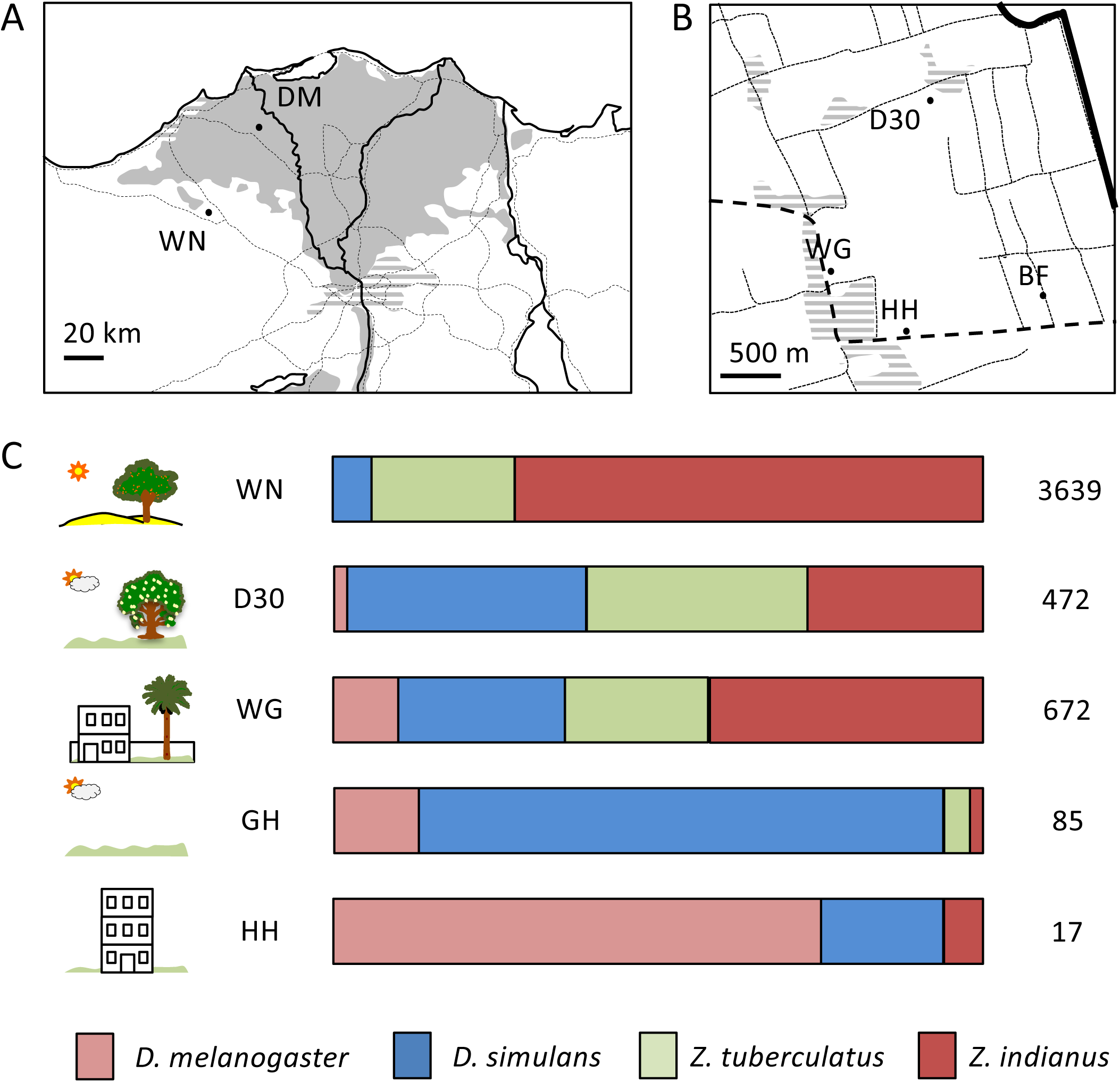
Drosophilids distribution in Egypt. A) Map of Northern Egypt showing the two collection sites: Wadi El Natrun (WN) in the desert (blank area) and Damanhur (DM) in the agrarian Nile Delta (shaded area). Transportation network and urban agglomerations are presented in dashed lines and striped areas, respectively. B) Microhabitats in DM including a guava orchard (D30), a house garden surrounded by date palm trees (WG), a house roof (HH) and a beet field with no fruit production (GH). C) Proportions of the four drosophilid species in WN and the four microhabitats of DM: *Drosophila melanogaster* (pink), *D. simulans* (blue), *Zaprionus tuberculatus* (green) and *Z. indianus* (red). The total number of collected individuals is given in front of each barplot.

For comparative phenotypic analyses between native and introduced populations, a few proportions of flies were kept alive in rearing bottles to establish mass cultures. For these analyses we used an introduced strain of *Z. tuberculatus* from D30. We also purchased a strain of *Z. indianus* from the Drosophila Species Stock Center (DSSC#50001-1031.00) that was collected from Alexandria, Egypt in 2006 from an agrarian environment similar to that of D30. For the native populations, we used two strains for both species that were collected from the island of Ngazidja (Grande Comore) from the Comoros archipelago in November 2018.

### Genome assembly

To investigate genetic diversity, we used two assemblies. For *Z. indianus*, we used the recently published assembly from the Drosophila 101 Genomes project GCA (Kim et al. 2021). This assembly has 1,078 scaffolds (total length 197 Mb) with L50 of 6.77 Mb. Benchmarking Universal Single-Copy Orthologs (BUSCO) v.5.0.0 (Manni et al. 2021) gave a score of 99.5% of the Dipteran conserved single-copy orthologs (diptera_odb10) with 0.5% of duplicated genes [C:99.5%[S:99.0%,D:0.5%],F:0.1%,M:0.4%,n:3285,E:3.0%].

For *Z. tuberculatus*, we assembled a new genome from 30 flies from the D30 mass culture using a combination of Oxford Nanopore Technology (ONT) long-read and Illumina short-read sequencing. For the former, DNA was extracted as above and fragments less than 10-kb were eliminated using the SRE XS kit from Circulomics (https://www.circulomics.com/, Baltimore, Maryland, USA). The ligation sequencing kit SQK-LSK109 (ONT) was then used to prepare the sample, which was purified and repaired with magnetic beads, and adapters (barcoding kits EXP-NBD104 and EXP- NBD114) were ligated on DNA fragments. The ONT raw data size was 1.86 Gb in 213,421 reads (mean read length 8.7 kb, longest read of 71.7 Mb), with an N50 of 11.7 kb. Phred scores ranged from 7 to 16.5, with a median of 12.66, as assessed by PycoQC (Leger and Leonardi 2019). Illumina paired-end sequencing from Congo (see above) was used for hybrid assembly. Short and long reads were then assembled using the MaSurCa software package ver. 4.0.3 with Cabog assembler (Zimin et al. 2013). The assembly has 12,819 scaffolds (total length 181 Mb) with an L50 of 185,263. Its completeness was estimated to 98.1% with 4.3% duplicated genes using BUSCO on the Dipteran dataset [C:98.1%[S:93.8%,D:4.3%],F:0.5%,M:1.4%,n:3285,E:2.8%].

### Population genomics analyses

For introduced populations, genomic DNA was extracted using standard protocol of Nucleobond AXG20 (Macherey Nagel 740544) with NucleoBond Buffer Set IV (Macherey Nagel 740604) from a pool of wild-caught females from one desert (WN) and one agrarian (D30) population for each *Zaprionus* species in Egypt (*N* = 30). Illumina paired-end short-read sequencing was performed by Novogene Company Limited as stated above. For native populations, genomic DNA was also extracted from pools of females from one population of *Z. indianus* collected from Soulou, Mayotte in 2017 (*N* = 12), and one population from the island of Nosy Bé, Madagascar collected in 2009 (*N* = 30) and provided courtesy of late Jean R. David. For *Z. tuberculatus*, three females were sampled from the F_1_ of 10, ethanol-preserved isofemale lines (*N* = 30) collected from Brazzaville, Congo and kindly provided by Joseph Vouidibio in 2005. Pools of native populations were sent in ethanol to End2end Genomics LLC, Davis, California for DNA extraction, PE library preparation using NEBNext Protocol and double-paired Illumina NovoSeq sequencing at 75x coverage.

Population genomics analyses were conducted as previously detailed in Ferreira et al. (2021). In summary, reads were mapped on the corresponding species assembly using minimap2. Minimap2-generated SAM files were converted to BAM format using samtools 1.9 software (Li et al. 2009) and after keeping reads with mapping quality of ≥ 20. The BAM files were then cleaned and sorted using Picard v.2.0.1 (http://broadinstitute.github.io/picard/). Popoolation 2 software package v.1.20162 (Kofler et al. 2011) was used to generate a synchronized mpileup file, with the argument --min-quality 20 to retain only bases with quality scores ≥ 20. Pools were genotyped after only retaining sites with a minimum and maximum depths 10 and 100. To estimate the number of segregating sites (*S*), within-population polymorphism (*π*) and between-populations genetic differentiation (*F_ST_*) from these synchronized files we used a series of customized perl scripts all available on the Figshare folder associated with this manuscript, with all analyses conducted on non-overlapping windows of 100 kb.

### Identification of orthologous sets of loci

We annotated both reference genomes using a single round of Maker software package ver. 2.31.10 (Campbell et al. 2014) and using as guide the protein-coding-genes from four *Drosophila* species, namely *D. melanogaster*, *D. ananassae*, *D. innubila* and *D. virilis* as in Bastide et al. (2024) . Annotation was carried after masking repeats by first using RepeatModeler v2.0.1 (Flynn et al. 2020) to identify repeat-rich regions and then RepeatMasker v4.0.9 (Tarailo-Graovac and Chen 2009) to mask these regions. The number of annotated protein-coding-genes were 11,947 and 13,489 for *Z. indianus* and *Z. tuberculatus*, respectively. BUSCO assessed the completeness of annotation of repeat-masked genomes to be 93.4% [C:93.4%[S:92.9%,D:0.5%],F:1.7%,M:4.9%] and 92.7% [C:92.7%[S:88.3%,D:4.4%],F:2.0%,M:5.3%] for *Z. indianus* and *Z. tuberculatus*, respectively. BLAST (Altschul et al. 1997) was used to identify orthologous genes between the two species, as well as between *Z. indianus* and the complete *D. melanogaster* transcriptome downloaded from FlyBase (Thurmond et al. 2019). Only the highest hit was retained in each run. This has resulted into 9,858 protein-coding-genes that are found in the two *Zaprionus* species. However, because some orthologous genes may be found in low recombining regions in both or one species, hence complicating comparative analyses of polymorphism and differentiation levels between the two species, only genes with the proportion of segregating sites (*S*) to the number of evaluated sites fell between >5% and <95% quantiles in both species were retained. 8,129 genes satisfied these conditions and were the basis of comparative analyses.

### Testing for potential parallel selection targets

To test for potentially parallel selection targets, we estimated *Population Branch Statistics (PBS)* (Yi et al. 2010) and *Population Branch Excess (PBE)* (Yassin et al. 2016) from *F_ST_* estimates of each single gene ± 2 kb, present in both species (see above). *PBE* is a standardized version of the Population Branch Statistic (*PBS*), which was proposed to identify population-specific allelic changes from *F_ST_* pairwise comparisons between three populations. The standardization measures how *PBS* at a locus, deviates from the genome-wide median *PBS* divided by the median *F_ST_* estimate from the two non-focal populations. Therefore, *PBE* is less sensitive than *PBS* to genome-wide variation in polymorphism levels. Unlike *F_ST_* measurements between two populations, *PBE* informs on which population changes took place compared to the allele frequencies in the outgroup. We measured *PBS* and *PBE* for both the WN (desert) population of both species, as well as for the Afrotropical population for each species. For a given gene, *PBS* and *PBE* for the desert population are calculated as:

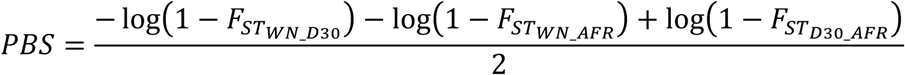

And

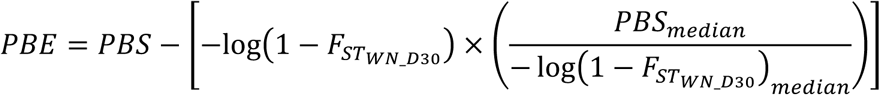

To avoid errors due to low or excess polymorphic genes, *PBS* and *PBE* were estimated after excluding genes with a proportion of segregating sites (*S*) <5% or >95% quantiles in either species. This has led to the retention of 8,129 orthologous genes between the two species. To test for genomic parallelism, we coded genes with *PBE* values <95% and ≥95% quantile as 0 and 1, respectively, and conducted an asymptomatic two-sample Fisher-Pitman permutation test using the oneway_test function in the coin package (Hothorn et al. 2006) in R v.4.3. (R Core Team 2023).

### Fruit preference assays

We adapted the population cages described above for multiple fruit choice assays. For each species, the same introduced (Egypt) and native (Comoros) populations were used. For each assay, 200 unsexed > 4-days old flies were introduced into a cage with a choice between 7 fruits, namely fig (dry), date (dry), banana (fresh), papaya (fresh), orange (fresh), pineapple (fresh) and guyava (juice). For each fruit, equal amounts (2 pieces, 1 cm x 1 cm each), were cut and placed in two empty culture vials.

The same disposition of the 14 vials were used in each cage. The plug of each vial (Flystuff droso-plugs®, Genesee Scientific) was pierced and a 200 µL pipette cone was placed through the hole. The cones were cut at the tip to allow the passage of a fly uniquely into the vial. The cages were placed in an incubator at 20°C with 50% relative humidity (RH) and left in the dark for 3 days. Flies trapped in each vial were then counted. Three replicates were done for each population. All statistical analyses were conducted under R.

### Desiccation resistance assays

We compared survival under desiccation conditions between the same introduced (Egypt) and native (Comoros) populations of both species. For each population, 64 >4-day old flies of the same sex were individually placed into a separate tube without food or water in two DAM2 Drosophila Activity Monitor (Trikinetics ®) devices. An agar mixture with no sucrose was placed into both ends of each tube. Each device simultaneously measures the locomotor activity of 32 individual flies through an infrared beam that crosses the tube at its midpoint and whose interruption by the fly movement is recorded. An agar mixture with no sucrose was placed into both ends of each tube. Each DAM2 device was placed in an incubator with a 12:12 hr light:dark cycle. As a control, a plate with water was added in the incubator, whereas for the desiccation assay, a plate with 150 g of silica gel was added. Relative humidity (RH) was controlled for each incubator. For each population, 64 flies of the same sex were tested under each condition (H_2_O and silica gel) under four temperatures: 18, 21, 25 and 28°C. Both males and females were tested for each population. The experiments lasted until no more movement was detected in the tubes or up to 1 week with the survival of each fly scored in minutes (min). Only flies which survived more than 120 min to reduce the effect of experimental handling. All statistical analyses were conducted under R.

## Results

### Distinct distributions of Z. indianus and Z. tuberculatus in Egypt

3,639 and 1,246 flies were collected from the desert locality (Wadi El Natrun, WN) and the agrarian region (Damanhour, DM), respectively. Taxonomic sorting of those flies unraveled *Z. indianus* prevalence in both environments (Figure 1C). *Z. indianus* predominated in the WN desert, accounting for nearly 72% of collected flies, followed by *Z. tuberculatus* (22%) and then *D. simulans* (6%), with *D. melanogaster* being completely absent. In the agrarian DM, *Z. indianus* still predominated by to a lesser extent, accounting for 41% of flies and followed by *D. melanogaster/simulans* (35%) and then *Z. tuberculatus* (24%). Closer investigation of those proportions revealed important differences (Figure 1C): the proportions of *Z. indianus* differed between 27%, 42%, 6% and 2% for D30, WG, HH and GH, respectively, whereas *Z. tuberculatus* differed between 34%, 22%, 0% and 4% for the same localities, respectively. Whereas both species are less abundant compared to *D. melanogaster/simulans* in highly urbanized or fields with no fruits, *Z. tuberculatus* is more frequent than *Z. indianus* in fruit crop orchards and open fields.

### Egyptian Zaprionus populations have low genomic differentiation

Within-population polymorphism levels, as measured in terms of nucleotide diversity (*π*), did not significantly differ between D30 and WN in each species, although it was slightly lower in *Z. tuberculatus* populations (*π* = 0.0176 ± 1.96 x 10^-4^ and 0.0153 ± 1.61 x 10^-4^ for *Z. indianus* and *Z. tuberculatus*, respectively, Student’s *t P* < 2.2 x 10^-16^; Supplementary Table S1 and Supplementary Table S2). Afrotropical populations of both species significantly differed from each other as well as each from conspecific Egyptian populations (*π* = 0.0151 ± 1.55 x 10^-4^ and 0.0224 ± 2.01 x 10^-4^ for Soulou and Nosy Bé *Z. indianus* populations, respectively, and 0.0181 ± 1.43 x 10^-4^ for *Z. tuberculatus*).

Remarkably, *π* was lowest in the small Afrotropical *Z. indianus* sample, which came from the island of Mayotte (*N* = 12), and highest for population collected from Nosy Bé (*N* = 30). Average genome-wide differentiation (*F_ST_*) between D30 and WN populations in Egypt was lower in *Z. indianus* than in *Z. tuberculatus* (*F_ST_* = 0.0209 ± 1.74 x 10^-4^ and 0.0261 ± 1.52 x 10^-4^, respectively; Student’s *t P* < 2.2x10^-16^; Supplementary Table S1 and Supplementary Table S2). Remarkably, the differentiation between Egyptian populations of both species was far lower from the differentiation between Egyptian and Afrotropical populations of both species (*F_ST_* = 0.1223 ± 9.93 x 10^-4^ and 0.1221 ± 9.59 x 10^-4^ for differentiation between D30 and Soulou or Nosy Bé *Z. indianus* populations, and 0.1398 ± 6.87 x 10^-4^ for *Z. tuberculatus*, respectively; Figure 2A and B) supporting a single introduction of each species in Egypt followed by local differentiation. However, average *F_ST_* between Egyptian and Afrotropical populations is higher in *Z. tuberculatus* (Student’s *t P* < 2.2x10^-16^). In summary, genetic differentiation is higher in *Z. tuberculatus* in agreement with the older expansion of this species in Egypt. For sake of comparison, *F_ST_* levels between introduced and native populations are presented at 8,129 genes ± 2 kb orthologous between the two species in Figure 2A and B.

**Figure 2.**
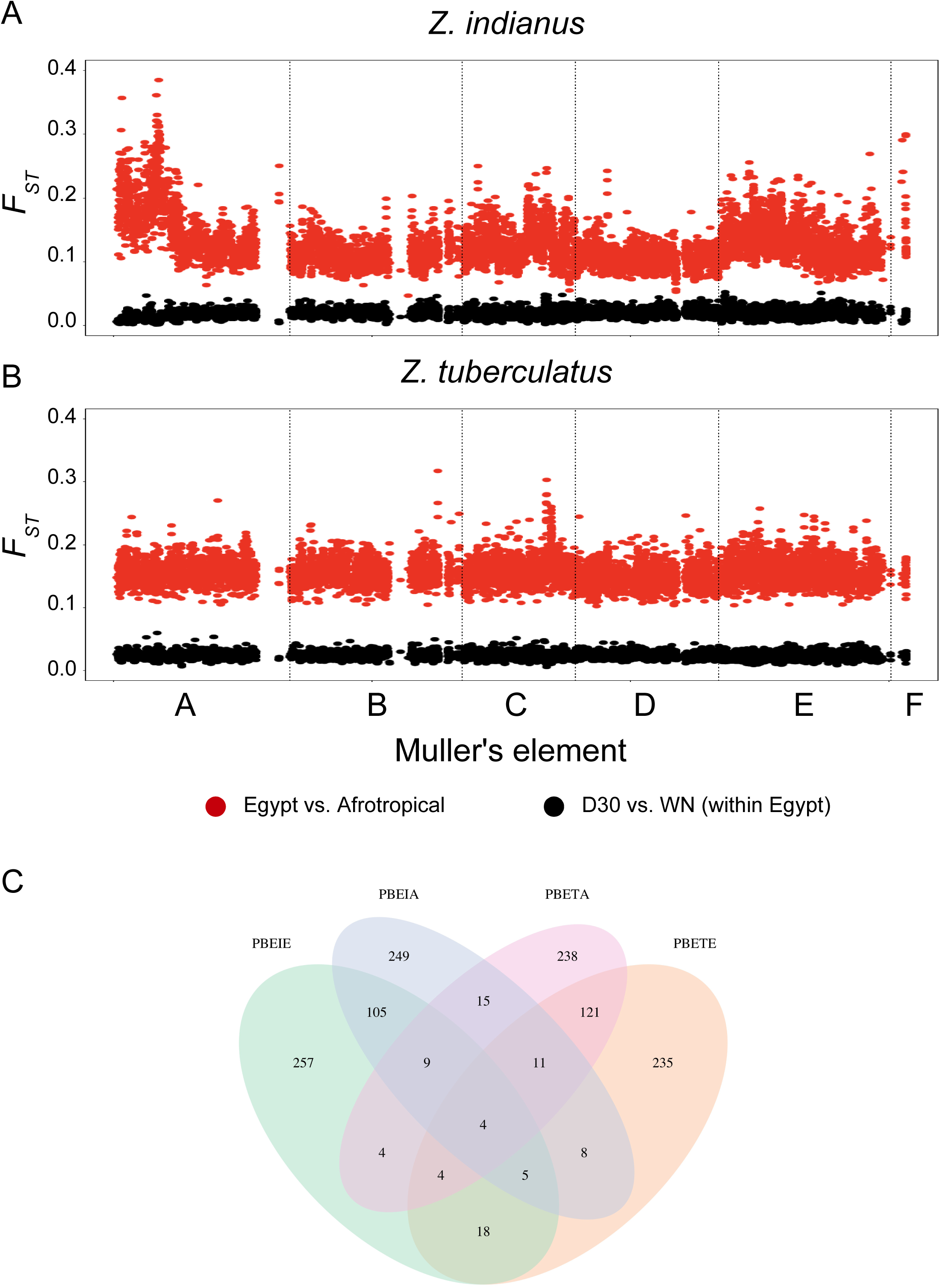
Low differentiation between Egyptian populations (black) relative to between Egyptian and Afrotropical populations (red) in A) *Z. indianus* and B) *Z. tuberculatus* at 8,129 orthologous genes ±2 kb. Genes are aligned according to the order of the *Z. indianus* reference genome with scaffold being ordered according to their sizes after tentative assignment of each scaffold to a Muller’s element according to the position of the *D. melanogaster* orthologs of their respective genes (A to F; dotted lines). C) Venn diagram’s showing shared orthologs with *PBE* values at the upper 5% quantiles in Egyptian and African *Z. indianus* (*PBEIE* and *PBEIA*, respectively) and *Z. tuberculatus* (*PBETE* and *PBETA*, respectively).

### Post- and pre-introductory parallelism of genetic differentiation at orthologous genes

In order to test for genetic parallelism, we compared pairwise *F_ST_* estimates at orthologous genes ± 2 kb between D30 and WN populations of the two species. We found significant correlation (*r* = 0.1407; *P* < 2.2 x 10^-16^). Similarly, *F_ST_* estimates significantly correlated between Egyptian and Afrotropical populations of the two species (*r* = 0.1342; *P* < 2.2 x 10^-16^). Because *F_ST_* estimates do not inform on the directionality of the allelic frequency change, we also applied *PBS* (Yi et al. 2010) and its standardized form *PBE* (Yassin et al. 2016) to infer population-specific changes. For *PBE*, 31 out of 8,129 genes fell ≥95% quantile in the WN population of both species ; (Supplementary Table S3; Figure 2C). This number exceeds the expected ∼20 genes that could have been commonly drawn by random in both species (Asymptotic two-sample Fisher-Pitman permutation test Z = -2.506, *P* = 0.0122). Similarly, 39 genes fell ≥95% quantile in the African population of both species compared to the two Egyptian ones (Asymptotic two-sample Fisher-Pitman permutation test Z = -4.376, *P* = 1.21 x 10^-5^; Supplementary Table S3; Figure 2C). These results suggest that selection has likely targeted the same set of genes pre- and post-introduction in Egypt.

### Egyptian Z. tuberculatus has distinct fruit attractions

When given simultaneous multiple choices between seven fruits, the two species showed distinct preferences (χ^2^ between-species = 907.79, df = 6, *P* = 7.789 x 10^-193^). On average, *Z. tuberculatus* had the strongest preference for figs whereas *Z. indianus* had the strongest preference for pineapple, regardless to the population of origin (Figure 3).

**Figure 3.**
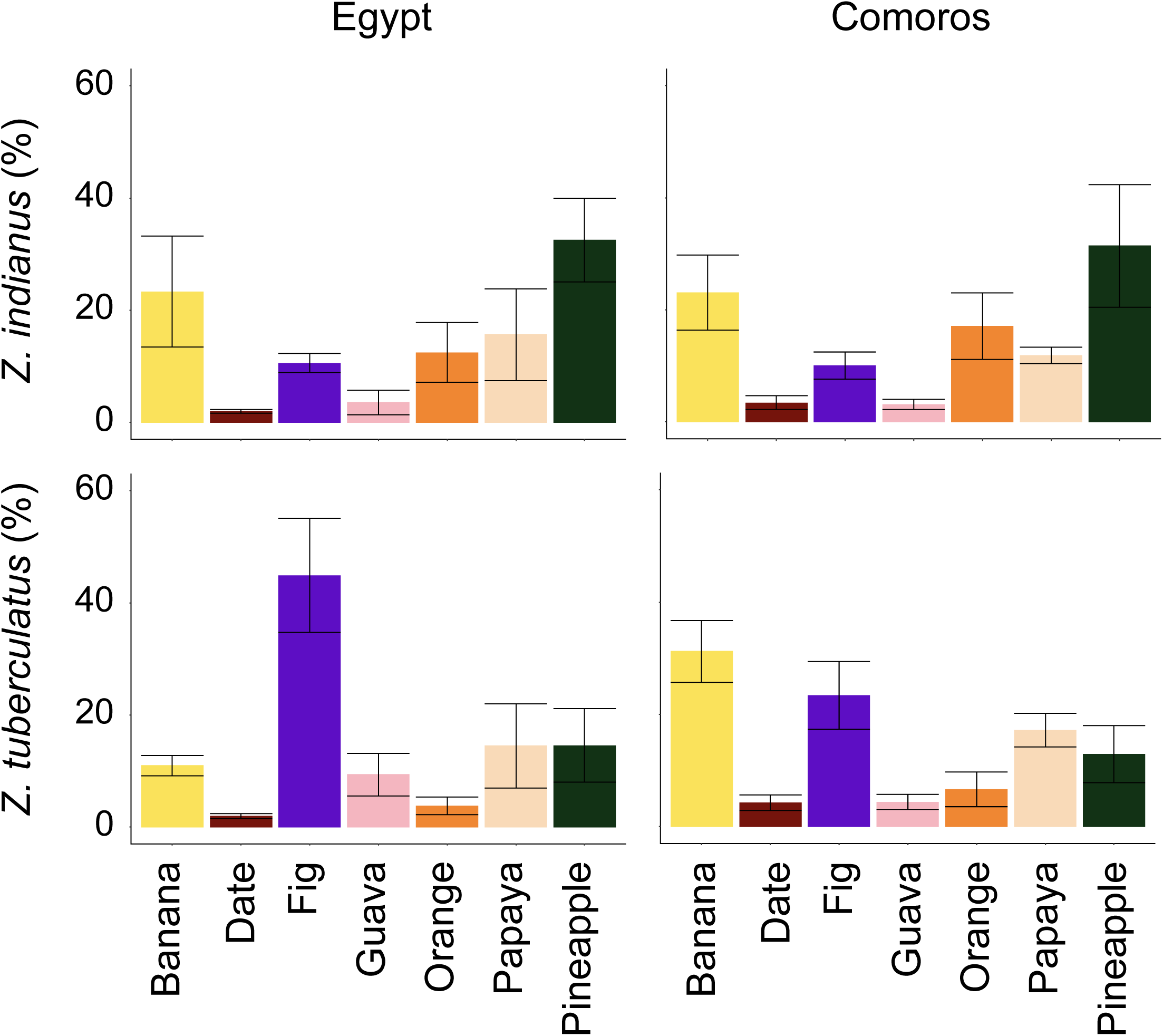
Fruit preference in introduced (Egypt) and native (Comoros) populations of *Z. indianus* and *Z. tuberculatus*. Bar heights reflect the frequency of captured flies (*N* = 200) in vials with each of the seven fruits, and lines indicate standard errors over 3 replicates.

However, when the Egyptian population was compared to the Afrotropical population in each species, *Z. tuberculatus* showed stronger difference (χ^2^ between-populations = 226.03, df = 6, *P* = 5.390 x 10^-46^) compared to *Z. indianus* (χ^2^ between-populations = 35.315, df = 6, *P* = 3.745 x 10^-6^). In particular, more than 50% of Egyptian *Z. tuberculatus* flies were attracted to figs more than any other fruits and to a lesser degree to guava and banana (∼10%). The Comoran population of the same species were nearly equivalently attracter to both figs and banana (∼25% each) and to a lesser degree to pineapple and papaya (∼10%). For *Z. indianus*, both Egyptian and Comoran populations showed strong attraction to pineapple (>30%), followed by banana and orange. Only a few proportions of *Z. indianus* flies were attracted to figs, despite the species vernacular name “the African fig fly”.

### Egyptian Z. tuberculatus has higher survival under cold and arid conditions

Multifactorial ANOVA indicated that all factors, namely species, population, temperature and treatment (H_2_O vs. silica gel) affected the survival of the flies in the desiccation experiments (*P* < 2.20 x 10^-16^ for all factors except for species, for which *P* = 9.84 x 10^-7^; Supplementary Table S4; Figure 4). On average, females lived longer than males, *Z. indianus* longer than *Z. tuberculatus*, Egyptian longer than Comoran, with water longer than with silica gel, and there was a decreasing trend of survival with increasing temperature. The relatively lower effect of the species is mainly due to an opposite trend in the population x species interactions (*P* < 2.20 x 10^-16^ in the multifactorial ANOVA). In Egypt, *Z. tuberculatus* survives longer especially for females under low temperatures, whereas in Comoros, *Z. indianus* lives longer. The difference between the temperate (Egyptian) and tropical (Comoran) populations was greater in *Z. tuberculatus* (1.66-fold, Student’s *t P* < 2.20 x 10^-16^) than in *Z. indianus* (1.19-fold, Student’s *t P* = 3.19 x 10^-10^), in agreement with the higher genomic and host preference divergence within *Z. tuberculatus*.

**Figure 4.**
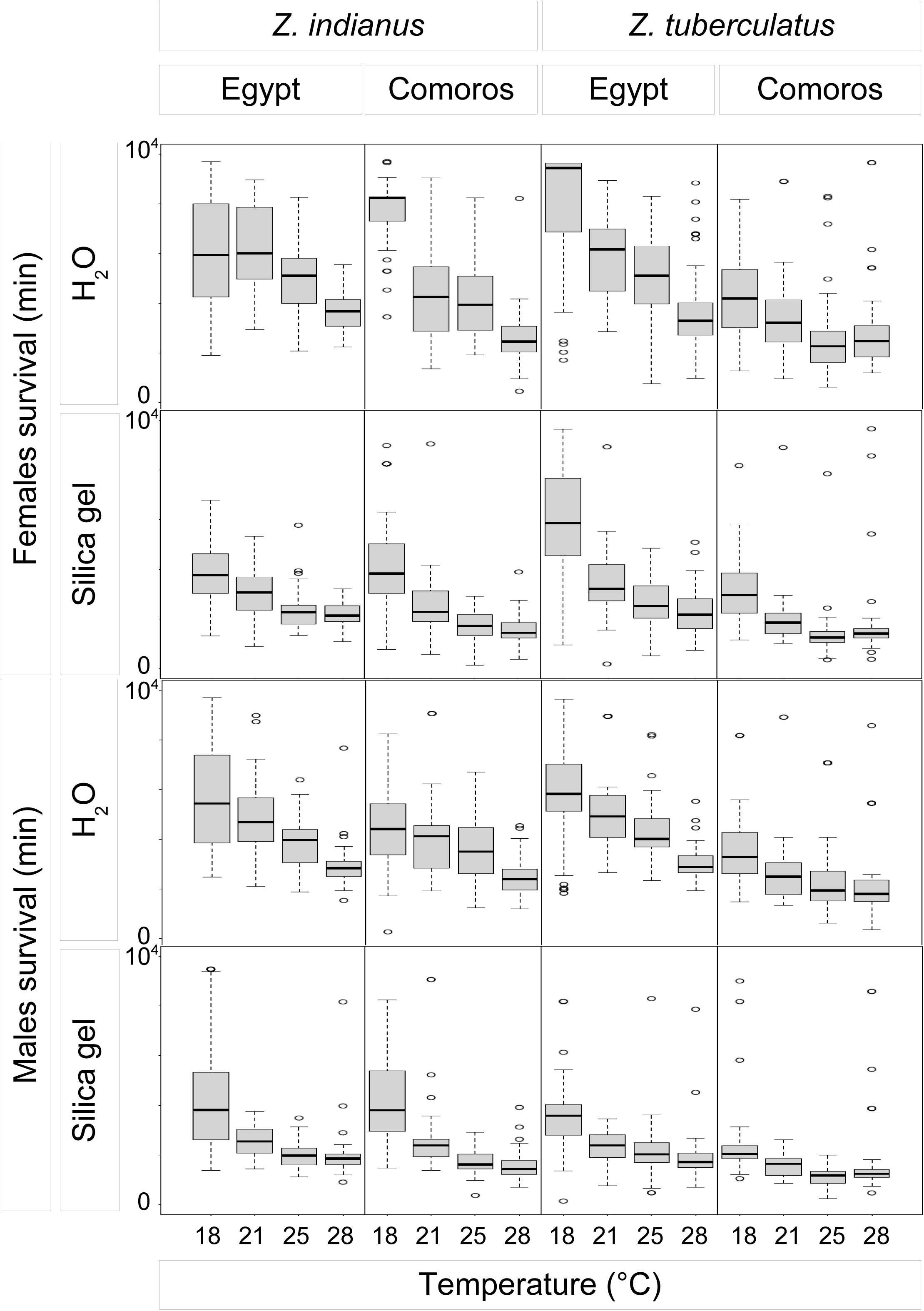
Survival plots in introduced (Egypt) and native (Comoros) populations of *Z. indianus* and *Z. tuberculatus* under controlled (H_2_O) and desiccation (SG: silica gel) conditions and under four temperatures.

## Discussion

The Egyptian drosophilid fauna is predominated by four species, all of Afrotropical origin. However, the temporal dynamics of the four species has been characterized by rapid turnovers. In the 1950s, *D. melanogaster* prevailed over *D. simulans* in most agrarian environments (Dobzhansky et al. 1957) but the pattern has been reversed in less than 10 years (Tantawy et al. 1970). After the collection of a few individuals of *Zaprionus tuberculatus* in Egypt in 1980 (Chassagnard and Kraaijeveld 1991), this species became the predominant figure in the early 2000s, when the fourth Afrotropical invader, *Z. indianus*, was detected for the first time (Yassin and Abou-Youssef 2004). Our study shows that, nearly 20 years later, *Z. indianus* became the predominant species, in both agrarian and desert environments. The order of overall prevalence of each species strikingly reflects the order by which they have been introduced in Egypt.

During the last 70 years, temperature increased and rainfall decreased in Egypt (Nashwan et al. 2019), which might have influenced the order of invasion and the outcome of the competition between the four species. Temperature indeed affects the outcome of laboratory competition between *D. melanogaster* and *D. simulans* (Tantawy and Soliman 1967; Montchamp-Moreau 1983). However, we still lack such experimental studies involving *Zaprionus* species, either in pairs or in groups. In the United States, competition with *Z. indianus* reduces *D. melanogaster* abundance by 10-53% during the summer, with a deep impact on the latter’s evolutionary trajectories in the fall when the two species no longer co-exist (Grainger et al. 2021). In Egypt, on the other hand, the four species co-exist during the four seasons (Elias 2005) but our results show that they have distinct microhabitat preferences and overlaps. *D. melanogaster* and *Z. indianus* are more adapted to hot weather than their congeneric species, *D. simulans* and *Z. tuberculatus*, respectively (David et al. 2004; Comeault et al. 2020). On the other hand, *D. melanogaster* and *Z. tuberculatus* are more resistant to desiccation (David et al. 2004; Kellermann et al. 2012; this study). Thus, complex interactions, that future laboratory experiments should investigate, may shape the competitive dynamics between the four species in Egypt.

Parallelism in genetic changes is more expected with higher degrees of genetic relatedness and similarities in the selective pressures (Conte et al. 2012; Stuart et al. 2017). Indeed, genomic parallelism between native and introduced populations of *D. melanogaster* and *D. simulans*, which have diverged 2 myr ago, was observed in response to latitudinal gradients (Machado et al. 2016; Sedghifar et al. 2016) and contrasting microhabitats (Kang et al. 2019). The two *Zaprionus* species studied here are far less related than the two *Drosophila* congeners (Yassin et al. 2008a; Suvorov et al. 2021).

Nonetheless, we were still able to detect a signal of parallel genome-wide differentiation between post-introduction desert and agrarian populations, as well as between introduced and native populations. Indeed, among the 31 most commonly- differentiating desert-specific genes four associate with thermal acclimation or stress in *D. melanogaster*, such as *Rpt4*, *Send1*, *CG15093* and *CG12848* (MacMillan et al. 2016; Gruntenko et al. 2021) and one, *Ir21a*, is involved in thermosensation (Knecht et al. 2016; Ni et al. 2016). At the pre-introductory level, we found two commonly- differentiated genes, *Achl* and *Cyp4p2*, to associate with circadian clock and sleep (Thimgan et al. 2015; Li et al. 2017), two phenotypes that can significantly vary between tropical and temperate environments.

Whereas our findings suggest that a small set of genes likely underlies invasion success even among distant lineages, we take this promising result with caution since we recognize some limitations in our population genomic analyses. For example, there is no chromosome-scale assembly for either *Zaprionus* species, and we did not assemble highly contiguous genomes from Egyptian flies. Such assemblies would have helped illustrating the boundaries of chromosomal inversions, evidence for which have already been shown for both species (Gupta and Kumar 1987; Elias 2005; Ananina et al. 2007; Yassin et al. 2009). Because inversions affect polymorphism levels and consequently differentiation statistics (e.g., *F_ST_*, *PBE*), presence of distinct inversions in each species can diminish the power of detection of common selection targets falling outside these inversions (Ferreira et al. 2023). We also only used Pool-Seq data for both Egyptian and Afrotropical flies. Genetic differentiation estimates can be more biased, especially by the variation of the mapping depth, in Pool-Seq data (Hivert et al. 2018). However, *PBE* which corrects for polymorphism variation across loci provides similar outliers in Pool-Seq and individual-sequenced (Ind-Seq) data (Ferreira et al. 2023), and this observation needs to be reconfirmed in future population genomics analyses of *Zaprionus* flies using Ind-Seq. Recent studies have indeed published sequences for a few Afrotropical populations for both species using Ind-Seq approaches (Comeault et al. 2020, 2021).

Future Ind-Seq analyses of flies from Egypt and additional African populations should help better illustrating their differences and infer a proper demographic model for the history of expansion. Besides, our populations were sampled on a single collection time (November 2019). Future comparisons of seasonal variation can greatly elucidate common targets of temporal selection in both species (e.g., Behrman et al. 2015; Machado et al. 2016, 2021).

Despite these limitations, our analyses support a consistent pattern: Egyptian *Z. tuberculatus* are more genetically, behaviorally and physiologically divergent from their Afrotropical populations. *Z. tuberculatus* may therefore be more adapted to the temperate environment than their sympatric *Z. indianus* that has invaded the Palearctic region in more recent times. *Z. tuberculatus* is currently invading Southern Europe (Georges et al. 2024) and South America (Cavalcanti et al. 2022). These continents also show several contrasting environments. Understanding the phenotypic divergence among multiple Palearctic and Afrotropical populations of this species is essential to identify the origin of the propagules and retrace post-introductory adaptations in invasive populations on the three continents. The comparative analysis of the Egyptian populations of the two species presented here is therefore a first significant step towards this goal.

## Acknowledgments

This work was funded by a PHC IMHOTEP program from the Egyptian Academy of Scientific Research and Technology (ASRT) and the French ministries of European and Foreign Affairs (MEAE) and High Education, Research and Innovation (MESRI) to A.M.E.S. and A.Y., and by the Agence Nationale de la Recherche (ANR) grant (ANR-18- CE02-0008) to A.Y. The authors thank Mr. M. Saïd El-Wakil for his help facilitating sample collection.

## Data availability statement

Population genomics and R scripts and data associated with this study are available at https://figshare.com/s/a7bc5d704977f9ba7f3b. Read sequences are deposited on NCBI Sequence Read Archive (SRA) associated to the Bioproject (PRJNA1164717).

## Conflict of interests

The authors declare no conflict of interests.

## Supplementary Tables

**Supplementary Table S1** – Nucleotide diversity (*π*) and genetic differentiation (*F_ST_*) in 100-kb long windows in Egyptian and African *Z. indianus*.

**Supplementary Table S2** – Nucleotide diversity (*π*) and genetic differentiation (*F_ST_*) in 100-kb long windows in Egyptian and African *Z. tuberculatus*.

**Supplementary Table S3** – Genetic differentiation (*F_ST_*) and *Population Branch Excess (PBE)* in 8,129 orthologous genes between *Z. indianus* and *Z. tuberculatus*.

**Supplementary Table S4** – Multifacotrial Analysis of Variance (ANOVA) for the effects of species, population, treatment, temperature and their interactions in the desiccation resistance experiments.

## Notes

https://figshare.com/s/a7bc5d704977f9ba7f3b

### Competing Interest Statement

The authors have declared no competing interest.

